# Langerhans islets induce anti-tumor immunity at the expense of glycemic control and predict chemotherapy response in pancreatic cancer

**DOI:** 10.1101/2021.03.13.435100

**Authors:** Azaz Ahmed, Pornpimol Charoentong, Rosa Klotz, Sophia Köhler, Meggy Suarez-Carmona, Nektarios A. Valous, Dyke Ferber, Mathias Heikenwälder, Nathalia Giese, Thilo Hackert, Fee Klupp, Martin Schneider, Thomas Schmidt, Christoph Springfeld, Yakup Tanriver, Christine S. Falk, Laurence Zitvogel, Inka Zörnig, Dirk Jäger, Niels Halama

## Abstract

Induction of anti-tumor immunity in pancreatic ductal adenocarcinoma (PDA) is an unresolved challenge. Systematic investigation of the microenvironment of primary pancreatic tumors revealed a role of endocrine Langerhans islets in the coordination of immune activation. We found that intratumoral β-cells, regulated via STAT3, secrete C-C motif chemokine ligand 27 (CCL27) and thereby promote a T_H_1 phenotype in the microenvironment resulting in an enhanced T cell infiltration and prolonged patient survival. The local effect can be abrogated in a patient-based human explant model by inhibition of the CCL27 receptor CCR10. This defense mechanism is paralleled by an impaired metabolic function of Langerhans islets with reduced insulin levels resulting in a dysregulation of glycemic control in patients. Based on these findings, screening of PDA cases (n= 2264) led to the identification of type 2 diabetes mellitus (T2DM) and extractable glycated haemoglobin (HbA1c) levels as response markers for neoadjuvant chemotherapy with fluorouracil, leucovorin, irinotecan and oxaliplatin (FOLFIRINOX). Collectively these data provide insights into the interconnection of T2DM and PDA, and link declining glycemic control to therapeutic efficacy, which can be utilized as a tool for clinical decision-making and improve patient management.

## INTRODUCTION

Pancreatic ductal adenocarcinoma (PDA) is the seventh-leading cause of cancer-related death worldwide, with a 5-year survival of only 9% and the highest incidence-to-mortality ratio of any solid cancer^1^. Over the past decade, not only has the involvement of immune system in malignancies emerged as a critical hallmark of cancer^2^, but immunotherapy has also profoundly changed cancer treatment by improving survival of patients in multiple solid tumors. A major predictor of clinical response to immunotherapy is the extent of intratumoral T cell infiltration. However, such “hot” tumors (high T cell infiltration) stand in contrast to “cold” tumors (low T cell infiltration) like PDA, which remains mostly refractory to immunotherapeutic treatment regimens^3^. A main feature of the tumor tissue in PDA is its desmoplastic microenvironment, in which immune cells make up nearly 50% of the cellular components but only few of them are anti-tumor effector cells such as CD8^+^ T lymphocytes^4^. Human studies have shown that paucity of T cells is common in PDA, but a subset of primary tumors do exhibit moderate infiltration of CD4^+^ or CD8^+^ T cells, and their sheer presence correlates with expression of cytotoxicity genes and overall survival^5-8^. Induction of such an anti-tumor immunity can be induced, for example, by chemotherapy with fluorouracil, leucovorin, irinotecan and oxaliplatin (FOLFIRINOX) by increasing cytotoxic T cell densities. To further improve patient outcomes, we need to understand the role of the components defining the tumor microenvironment and identify critical immunomodulatory mechanisms^9-11^.

In this regard, little is known about the role of endocrine Langerhans islets in shaping the tumoral immune landscape. Previous studies describe multiple proinflammatory cytokines originating from Langerhans islets in metabolic disorders such as obesity and diabetes^12,13^. Also, Langerhans islets are mainly responsible for chemokine-mediated inflammation and T cell infiltration in type 1 diabetes mellitus^14^ and evidence suggests that this patient cohort is protected against PDA^15^. But what phenotype do Langerhans islets exhibit in the immune contexture of PDA? And is it clinically relevant? In addition, an impaired glycemic control is a common finding and several studies have shown a high incidence of type 2 diabetes mellitus (T2DM) in PDA patients^16^ but it remains unclear whether T2DM is a risk factor or early marker of the disease^17-19^.

This study aimed to systematically assess the tumor microenvironment and subsequently decipher immunologic interactions in PDA patients. This led to the in-depth investigation of anti-tumoral immunomodulatory capacities of Langerhans islets, their consequences for glycemic control and their impact on clinical outcomes, in order to tailor therapeutic strategies by exploiting metabolic parameters as a stratification marker.

## MATERIAL AND METHODS

### Patient samples and clinical analyses

All patients underwent planned surgery at the department for General, Visceral and Transplantation Surgery at the University Hospital Heidelberg. The study was approved by the local ethics committee (301/2001 and S-457/2019). Written informed consent was obtained from all patients. Tissue samples of 61 PDA patients were used for immunohistochemistry and cytokine analyses. 9 fresh PDA tissues on ice were directly collected from the operating room from patients who underwent surgery in 2019 and 2020. Survival analyses were performed on a set of 51 patients with PDA. Clinical information about diagnosis of T2DM, HbA1c levels and neoadjuvant treatment with FOLFIRNOX was collected in a cohort of 2264 patients, which is the total number of patients which was planned for pancreatoduodenectomy from 2016-2020 in the University Hospital Heidelberg. 122 patients were included in the final analysis. All patient data and samples were collected in a prospective database and analyzed retrospectively. All patient data was pseudonymized.

### Patient-based organotypic tumor explant model

The organotypic tumor explant model was set up as previously described^20^. In brief, resected specimens were transferred on ice and under sterile conditions from the operating room to the laboratory. The timeframe was less than thirty minutes upon resection. The resected specimen was immediately processed into serial explant-suitable thin tissue sections (size approximately 6 x 6 x 1 mm). From each patient, one or more samples were directly frozen as well as formalin-fixed and embedded in paraffin for reference purposes. Tissue culture was performed by placing two explants with 1 ml of Minimum Essential Medium with 1% GlutaMAX supplement (Gibco, USA; 41090-036) per well on a 24-well plate in an incubator at 37°C and 5% CO_2_. The explants were treated with a small molecule inhibitor of CCR10 (BI-6901) from (Boehringer Ingelheim, Germany). The CCR10 antagonist dosage per well was 0.1 mg/ml. Along with untreated explants (reference), the treated specimens were harvested after 24 hours and cryo-preserved using cryo embedding medium (Medite, USA; 41-3011-00) as well as formalin-fixed and embedded in paraffin.

### Cell Culture of human pancreatic Langerhans islets

Human pancreatic Langerhans islet cells were purchased from (Celprogen, USA; 35002-04). The cells were seeded at 80% of confluence and plated at 10^5^ per ml density. The cells were maintained in the appropriate size, extra-cellular matrix precoated flasks (Celprogen, USA; 35002-04-T75) and cultured in complete growth medium with serum (Celprogen, USA; M35002-04S). Medium was changed every 24 hours. When reached 65-75% of confluence, cells were passaged with 0,05% Trypsin-EDTA (Gibco, USA; 25300-054) and subcultured for expansion. Langerhans islet cells from passages 3 to 5 were used for the experiments.

### Immunohistochemistry on human tissue samples

PDA tissue samples were collected freshly from the operating room, fixed in 4% phosphate buffered formaldehyde (ROTI Histofix, Roth, Germany; P087), placed in ethanol and embedded in paraffin. Tissue samples were sliced in 3 μm thick sections. All immunohistochemical stainings were performed on a BOND-MAX (Leica, Germany). FFPE tissues were deparaffinized and rehydrated (BOND Dewax Solution, Leica, Germany; AR9222). After heat-induced epitope retrieval (HIER) at 100°C (BOND Epitope Retrieval Solution 1 or 2, Leica, Germany; HIER 1 AR9961, HIER 2 AR9640), endogenous peroxidase activity was blocked by incubation with 3% peroxide block for 20 minutes (BOND Polymer Refine Detection System, Leica, Germany; DS9800). The sections were blocked with 10% normal goat serum (Vector, USA; S-1000-20). The primary antibodies were applied at room temperature for 30 minutes: CD3 (1:100, HIER 1, rabbit monoclonal, clone SP7, Abcam, UK; ab16669), CD4 (1:100, HIER 1, mouse monoclonal, clone 4B12, Leica, Germany; CD4-368-L-CE-H), CD8 (1:50, HIER 1, mouse monoclonal, clone 4B11, Leica, Germany, CD8-4B11-L-CE), CD20 (1:100, HIER 1, mouse monoclonal, clone L26, Leica, Germany; CD20-L26-L-CE), CD163 (1:500, HIER 2, rabbit monoclonal, clone EPR19518, Abcam, UK; ab182422), NKp46 (1:175, HIER 1, monoclonal mouse, clone 195314, R&D Systems, USA; MAB1850-500), FoxP3 (1:100, HIER 2, mouse monoclonal, clone 236A/E7, Thermo Fisher Scientific, Germany; 14-4777), CCL27 (1:200, HIER 1, mouse monoclonal, clone 124308, R&D Systems, USA; MAB376), CCR10 (1:100, HIER 1, rabbit polyclonal, Novus Biologicals, USA; NB100-56319), CA19-9 (1:1000, HIER 1, mouse monoclonal, clone 121SLE, Abcam, UK; ab3982), insulin (1:500000, rabbit monoclonal, clone EPR17359, Abcam, UK; ab181547), Phospho-Stat3 (Ser727) (1:200, HIER 2, mouse monoclonal, Cell Signaling Technology, USA; 9136S). The slides were incubated with a secondary antibody (rabbit anti-mouse IgG, BOND Polymer Refine Detection System, Leica, Germany, DS9800) for 8 minutes at room temperature. Signal amplification was performed by incubation with horse radish peroxidase and coupled to dextrane molecues in large numbers, for 8 minutes at room temperature (Poly-HRP-mouse anti-rabbit IgG, BOND Polymer Refine Detection System, Leica, Germany; DS9800). A color reaction with 3,3-di-amino-benzidine (DAB chromogen, BOND Polymer Refine Detection System; DS9800) was utilized to detect the antigen. Counterstaining was performed with haematoxylin (BOND Polymer Refine Detection System, Leica, Germany; DS9800) and the sections were mounted with Aquatex (Merck, Germany; 108562).

### Immunofluorescence on human tissue samples

Immunofluorescence double staining was performed on paraffin-embedded sections using fluorochrome-conjugated secondary antibodies. For the first primary antibodies CD3, CD4, CD8 and FoxP3 a red fluorescence Alexa Fluor 594 donkey anti-mouse IgG (Thermo Fisher Scientific, Germany; A-21203) was used, and for CCR10 a green fluorescence Alexa 488 goat anti-rabbit IgG (Thermo Fisher Scientific, Germany; A-11008). For the analysis of the first primary antibody CCL27 and Phospho-Stat3 (Ser727) a green Alexa Fluor 488 goat anti-mouse IgG (Thermo Fisher Scientific, Germany; A-11029) was used, and for insulin a red Alexa Fluor 594 donkey anti-rabbit IgG (Thermo Fisher Scientific, Germany; A-21207). The sections were stained according to antibody recommendation. After incubation of the first primary antibody overnight at 4°C, Alexa Fluor 488 (1:100 dilution) was applied for 60 minutes. The second primary antibody was applied for 180 minutes at room temperature and detected with Alexa Fluor 594 (1:100 dilution) for 60 minutes during sequential double staining. Mounting and staining for cell nuclei was performed using Vectashield with DAPI (1:10000, Vector, USA; H-1200). Images were taken with a Nanozoomer 2.0-HT slide scanner (Hamamatsu, Japan).

### Whole-slide immune cell quantification

For quantification analysis of immune cells, tissue sections were digitized with a Leica Aperio AT2 scanner (Leica, Germany). Whole-slide microscopic images of full tissue sections were automatically obtained (virtual microscopy). The slides were scanned at 40-fold magnification and further examined using an image analysis software (VIS software suite, Visiopharm, Denmark). Given regions of interests (tumor and stroma) were manually annotated and density and distribution of immune cells was analyzed semi-automatically, as reported previously^21-23^. A visual consistency check was performed on all evaluations.

### Laser capture microdissection

The tumoral and stromal compartment on PDA tissue sections was separated from each other by laser capture microdissection. This technique permits to isolate selected human cell populations from a section of complex tissue under direct microscopic visualization. The standard protocols of the inventers were used^24^. In brief, the tissue section was focused (20-fold magnification) and the tumoral and stromal area was manually separated using the Leica Laser Microdissection Software (Leica, Germany); the dissection was completed using a carbon dioxide laser pulse.

### Multiplex protein quantification of cytokines and metabolic hormones

Small pieces of dissected frozen pancreatic tumor tissues were collected and lysed (Bio-Plex Cell Lysis Kit, BioRad, USA; 171304011), followed by a repeated procedure of vortexing, freezing (−80°C for 10 minutes), thawing (on ice) and ultrasonic bathing (for 10 minutes). The supernatants were then centrifuged at 13.000 rpm for 20 minutes at 4°C. When supernatants of cell culture experiments were analyzed no lysis was performed. In all samples, the protein concentrations were determined (Pierce BCA Protein Assay, Thermo Fisher Scientific, Germany; 23227), the concentration then was adjusted to 200 μg/ml and cytokine as well as metabolic hormone concentrations were quantified by multiplex protein arrays, according to the instructions of the manufacturer (BioRad, USA). A two-laser array reader simultaneously quantifies all proteins of interest (cytokines/metabolic hormones) and concentrations are calculated with Bio-Plex Manager 4.1.1 based on a 5-parameter logistic plot regression formula. For cytokine quantification Bio-Plex Pro Human Cytokine Screening Panel 48-plex (BioRad, USA; 12007283), Bio-Plex Pro Human Cytokine ICAM-1 (BioRad, USA; 171B6009M) and Bio-Plex Pro Human Cytokine VCAM-1 (BioRad, USA; 171B6022M) were used. For the quantification of metabolic hormones Bio-Plex Pro Human Diabetes 10-plex Assay (BioRad, USA; 171A7001M) was used. Previous experimental insights showed the technical reproducibility of this protocol ^20^.

### Human Langerhans islets in tumoral microenvironment

Langerhans islets cells were placed in a 96-well plate (around 7500 cells/ per well) and cultured in a mixture of 150 μl volume of complete growth medium with serum (Celprogen, USA; M35002-04S) and 50 μl of supernatant of human PDA tissue explants. For reference purposes Langerhans islets cells were cultured in 200 μl of complete growth medium only. The supernatants were used to mimic a PDA tumor microenvironment and to investigate the behavior of intratumoral Langerhans islet cells. Supernatants of explants from five different PDA patients were used. The cells were treated for 48 hours. Afterwards, the total supernatants of the treated cells were collected and analyzed (Multiplex protein quantification of cytokines and metabolic hormones). For reference purposes the supernatant of undiluted human PDA tissue explants was also analyzed. The remaining cells were lysed (Bio-Plex Cell Lysis Kit, BioRad, USA; 171304011). In brief, the well-plate was placed on ice, the cells were rinsed with cell wash buffer and 120 μl of cell lysis buffer was added. The cells were incubated for 25 minutes on ice and during this time vortexed thoroughly every 5 minutes. After centrifugation (13.000 rpm for 30 minutes at 4°C), supernatant was collected and used for further analyses (signaling pathway analysis).

### Signaling pathway analysis

Protein concentrations of lysate samples were determined (Pierce BCA Protein Assay, Thermo Fisher Scientific, Germany; 23227), the concentration then was adjusted from 20 to 130 μg/ml. Afterwards, multiple phosphorylated and total proteins were simultaneously quantified in each well of 96-well plates, using a protein array system according to the instructions of the manufacturer (BioRad, USA). A dual-laser microplate reader system detects the fluorescence of the individual dyed beads and the signal intensity on the bead surface. The relative abundance of each target protein is reported as the ratio of fluorescence among the wells. In the present study, Bio-Plex Pro Cell Signaling MAPK Panel 9-plex (BioRad, USA; LQ00000S6KL81S) and Bio-Plex Pro Cell Signaling Akt Panel 8-plex (BioRad, USA; LQ00006JK0K0RR) were used for phosphorylated protein quantification and Bio-Plex Pro Total Akt (BioRad, USA, 171V60001M), Bio-Plex Pro Total ERK1/2 (BioRad, USA, 171V60003M), Bio-Plex Pro Total GSK-3β (BioRad, USA, 171V60004M), Bio-Plex Pro Total JNK (BioRad, USA, 171V60007M), Bio-Plex Pro Total MEK1 (BioRad, USA, 171V60008M), Bio-Plex Pro Total PTEN (BioRad, USA, 171V60016M), Bio-Plex Pro Total mTOR (BioRad, USA, 171V60015M), Bio-Plex Pro Total p38 MAPK (BioRad, USA, 171V60009M), Bio-Plex Pro Total p70 S6 Kinase (BioRad, USA, 171V60010M), Bio-Plex Pro Total Human GAPDH (BioRad, USA, 171V60019M), Bio-Plex Pro Total β-Actin (BioRad, USA, 171V60001M) were used for quantification of total proteins. The protein concentrations are calculated with Bio-Plex Manager 4.1.1 based on a 5-parameter logistic plot regression formula.

### CCL27 quantification from stained tissue sections

Samples were processed and stained automatically as described above. Whole-slide microscopic images were obtained and calibrated for proper comparability as described previously ^25^. Corresponding cytokine concentrations were used for reference and after calibration, tissues of interest were quantified for CCL27 positivity and concentrations were calculated based on reference curves.

### Statistics

Normality of the distributions was tested with Shapiro-Wilk test, and for normal distributed data the variance within each group of data was estimated and tested for equality between groups by a two-sided F-test. Cytokines which were undetectable in more than 10 percentage of samples were excluded. Correction for batch effects from three experiments was performed using the Python package *pycombat* (version 0.14) and validated by analyzing the mixture of samples within UMAP embeddings ^26^. For comparison of two patient groups, two-sided Student’s t test was used where stated, otherwise the non-parametric two-sided Wilcoxon-rank sum test was used. Based on the cytokine concentration data we reconstructed a T_H_1 cytokine-cytokine co-expression network. The edge weights of the network are based on the Pearson correlation coefficient between cytokine-cytokine (r ≥ 0, P < 0.05). Association of CCL27 concentrations with number of chemotherapy cycles was done using Kendall rank correlation. Resulting p-values were adjusted for multiple hypothesis testing according to the Benjamini and Hochberg method ^27^.

The overall survival time was defined using the latest information. For survival analysis, the patients were dichotomized based on cytokine concentration and cell density. The median cutpoints were determined to stratify patient into two groups (Hi and Lo). Kaplan Meier estimators of survival were used to visualize the survival curves. The log-rank test was used to compare overall survival between patients in different groups. P-values for the HiHi, HiLo, LoHi, and LoLo cytokine/cell combination analysis were corrected for multiple testing using the Benjamini-Hochberg method. All analyses were performed using the statistical software environment R (package *survival*). Stated percentages and quartiles include rounded numbers. Statistical analyses were carried out using R-4.0.2 and Python-3.6.4. Clustering and visualization were done with the software Genesis ^28^.

## RESULTS

### Intratumoral CCL27 expression is associated with improved overall survival

To investigate the quantities of intratumoral immune cells and corresponding cytokine levels in PDA, we systematically performed stainings for CD3^+^ T cells, CD8^+^ T cells, CD4^+^ T cells, FoxP3^+^ T cells, CD20^+^ B cells, NKp46^+^ natural killer cells and CD163^+^ tumor-associated macrophages and evaluated the expression profile of intratumoral cytokines in the resected tumor tissue of 51 PDA patients **(Table S1) (Figure S1A)**. These parameters were analyzed in relation to the clinical outcome of patients.

In tumor tissue, the only cytokine significantly associated with prolonged survival was the expression of C-C motif chemokine ligand 27 (CCL27) **(Figure 1A and 1B)**. Along with that, the tumor infiltration by CD8^+^ cytotoxic T lymphocytes and the intratumoral CD8^+^/CD163^+^-ratio was significantly associated with improved survival **(Figure 1B)**. Particularly, the combination of intratumoral CCL27 and CD8^+^ T cells proved to be a strong marker for favorable prognosis **(Figure 1B) (Table 1)**. On the other hand, intratumoral expression of IFN-γ was significantly associated with poorer survival in patients **(Figure 1A)**. The other intratumoral cytokines and immune cells tested were not significantly associated with overall survival **(Figure 1A)**. The association of clinical features (lymph node status, metastatic status, tumor grading) to overall survival confirmed previous reports **(Figure S1B)**.

**Figure 1.**
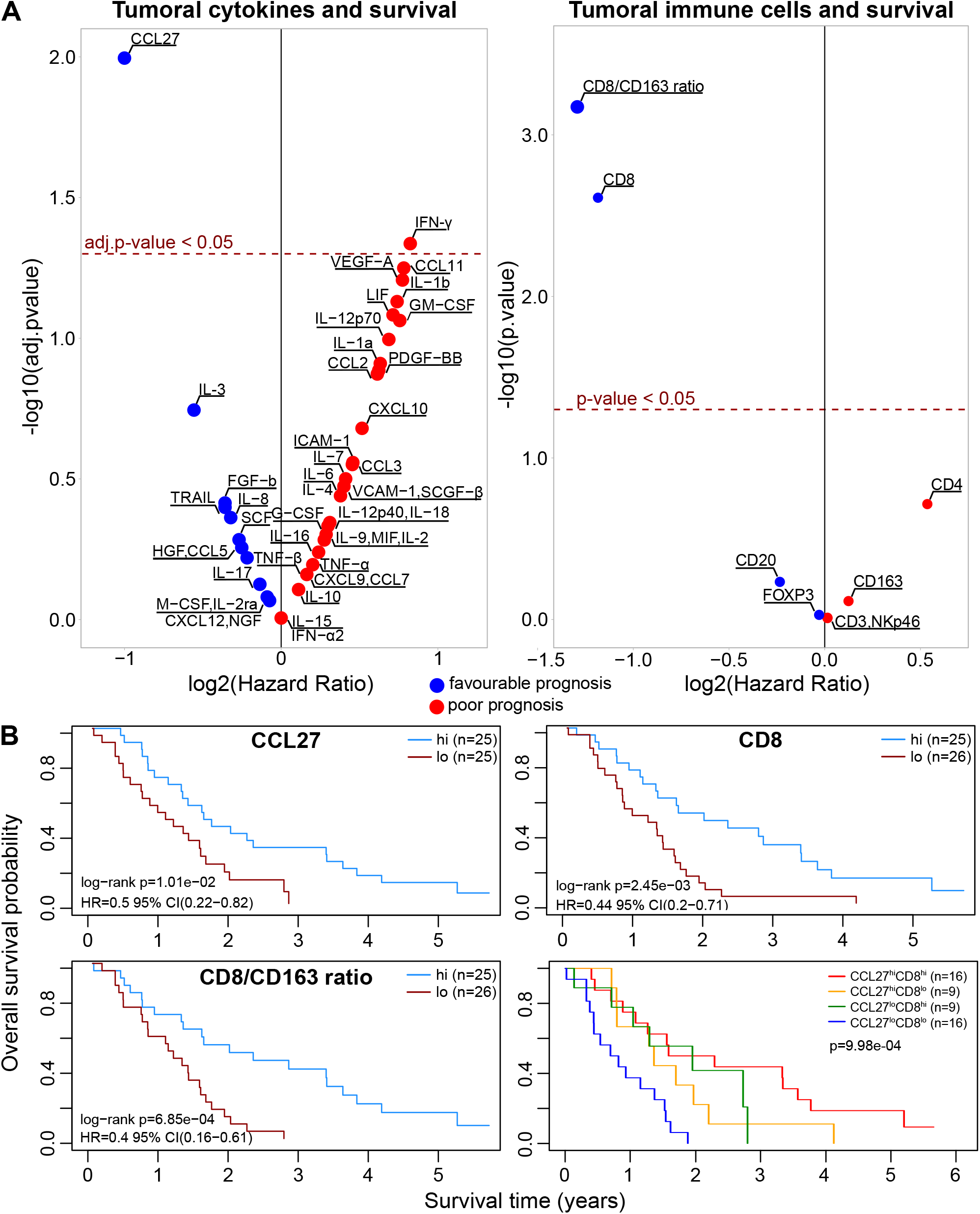
Intratumoral CCL27 expression is associated with improved overall survival **(A)** Volcano plot of statistical significance (y-axis) against log2(Hazard Ratio) (x-axis) for cytokines (left) and immune cells (right) showing favorable and poor prognostic markers. **(B)** Kaplan-Meier survival plots of patients with high (hi) versus low (lo) concentration of CCL27 (n=25/n=25), hi versus lo density of CD8+ T cells (n=25/n=26), hi versus lo CD8+/CD168+ ratio (n=25/n=26) and the combination of CCL27 (hi or lo) and CD8+ T cells (hi or lo). The median cutpoints were determined to stratify patients into hi and lo. Survival data were analyzed using the log-rank test. See also Figure S1.

**Table 1.**
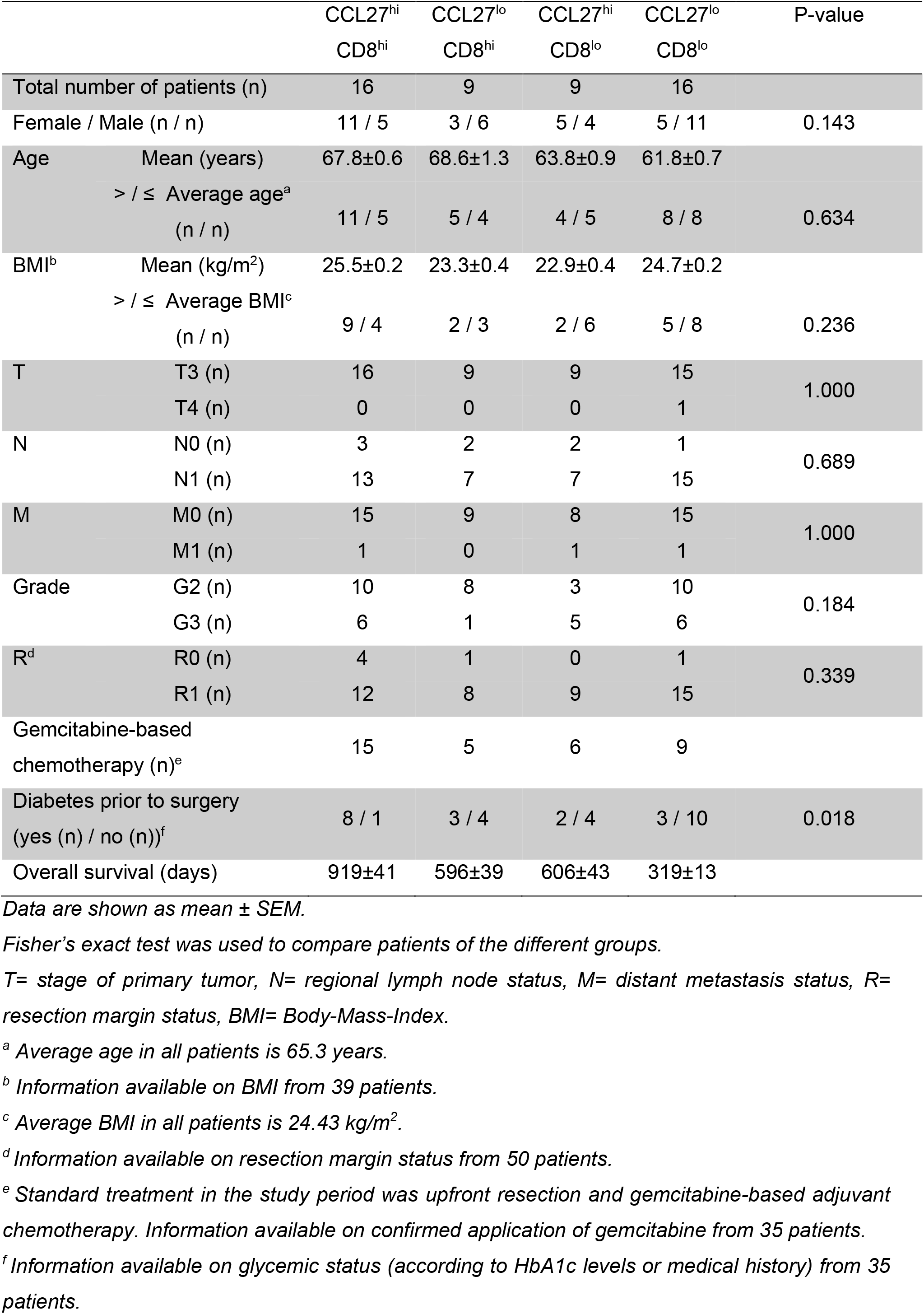
Subject characteristics

### β-cells within Langerhans islets are the source of CCL27 and its expression is correlated with a T_H_1-type cytokine program

To elucidate whether the intratumoral expression of CCL27 relates to an immunity-driven anti-tumor signature within the tissue of PDA patients, we performed cytokine-cytokine correlation analyses. By that, we observed a characteristic T_H_1-type cytokine signature correlating to the intratumoral presence of CCL27 **(Figure 2A and 2B)**.

**Figure 2.**
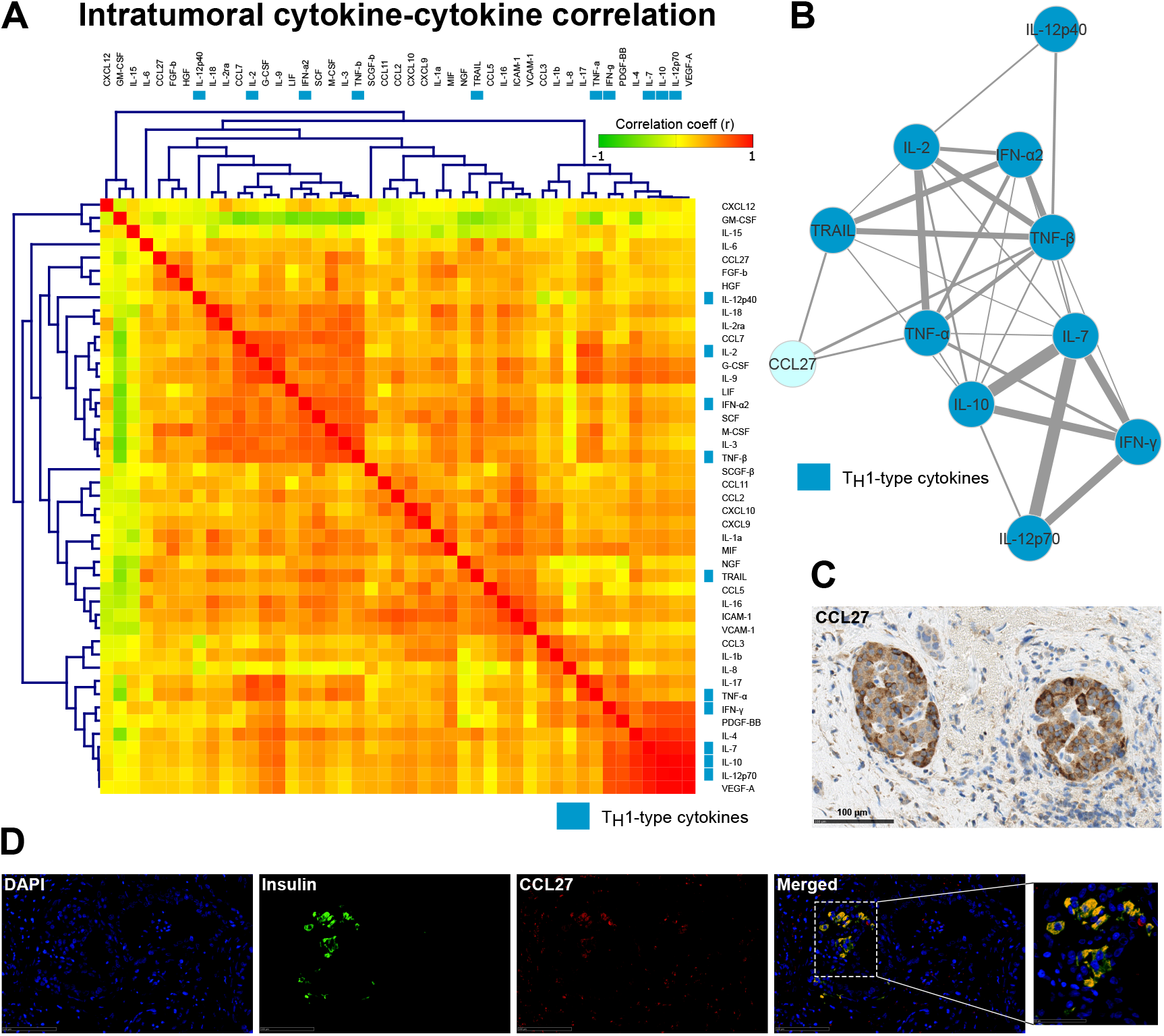
β-cells within Langerhans islets are the source of CCL27 and its expression is correlated with a TH1-type cytokine program **(A)** Heatmap showing the intratumoral cytokine-cytokine correlation from human PDA tissue samples (n=48). TH1-type cytokines are indicated in blue. **(B)** Network plot of intratumoral cytokine-cytokine correlations of TH1-type cytokines in human PDA tissue samples (n=48). The edge weights of the network are based on the Pearson correlation coefficient between cytokine-cytokine (r≥0, p<0.05). **(C)** Immunohistochemistry for CCL27 in a human PDA tissue sample. Scale bars, 100 μm. **(D)** Immunofluorescence for CCL27 and insulin (β-cell marker) in a human PDA tissue sample. Scale bars, 100 μm PDA= pancreatic ductal adenocarcinoma

Next, immunohistochemical and immunofluorescence stainings were performed on patient samples to identify the origin of CCL27 production. A staining for CCL27 revealed that islet-like structures within the tumor core are the main site of CCL27 expression **(Figure 2C)**. This observation prompted us to perform an immunofluorescence staining for the Langerhans islet marker insulin and CCL27, which corroborated that CCL27 is mainly expressed by β-cells within Langerhans islets **(Figure 2D)**.

### CCR10 is expressed on CD4^+^ FoxP3^-^ T cells and selective inhibition of CCR10 abrogates the T_H_1-type cytokine profile

Interaction of CCL27 with its only known receptor C-C chemokine receptor 10 (CCR10) is a key regulator for T-cell migration to the skin in inflammatory disorders^29^. And since our data revealed that tumor infiltration by cytotoxic T cells is independently prognostic favorable, we hypothesized that CCL27 might have similar immunomodulatory capacities in PDA. To identify CCR10-expressing cells in PDA, we performed systematic immunofluorescence stainings on patient samples for CCR10^+^ CD3^+^ T cells, CCR10^+^ CD4^+^ T cells, CCR10^+^ FoxP3^+^ T cells and CCR10^+^ CD8^+^ T cells. A strong co-positivity for CCR10 was observed on CD3^+^ and CD4^+^ T cells **(Figure 3A)**. FoxP3^+^ and CD8^+^ T cells showed negligible expression of CCR10 **(Figure 3A)**.

**Figure 3.**
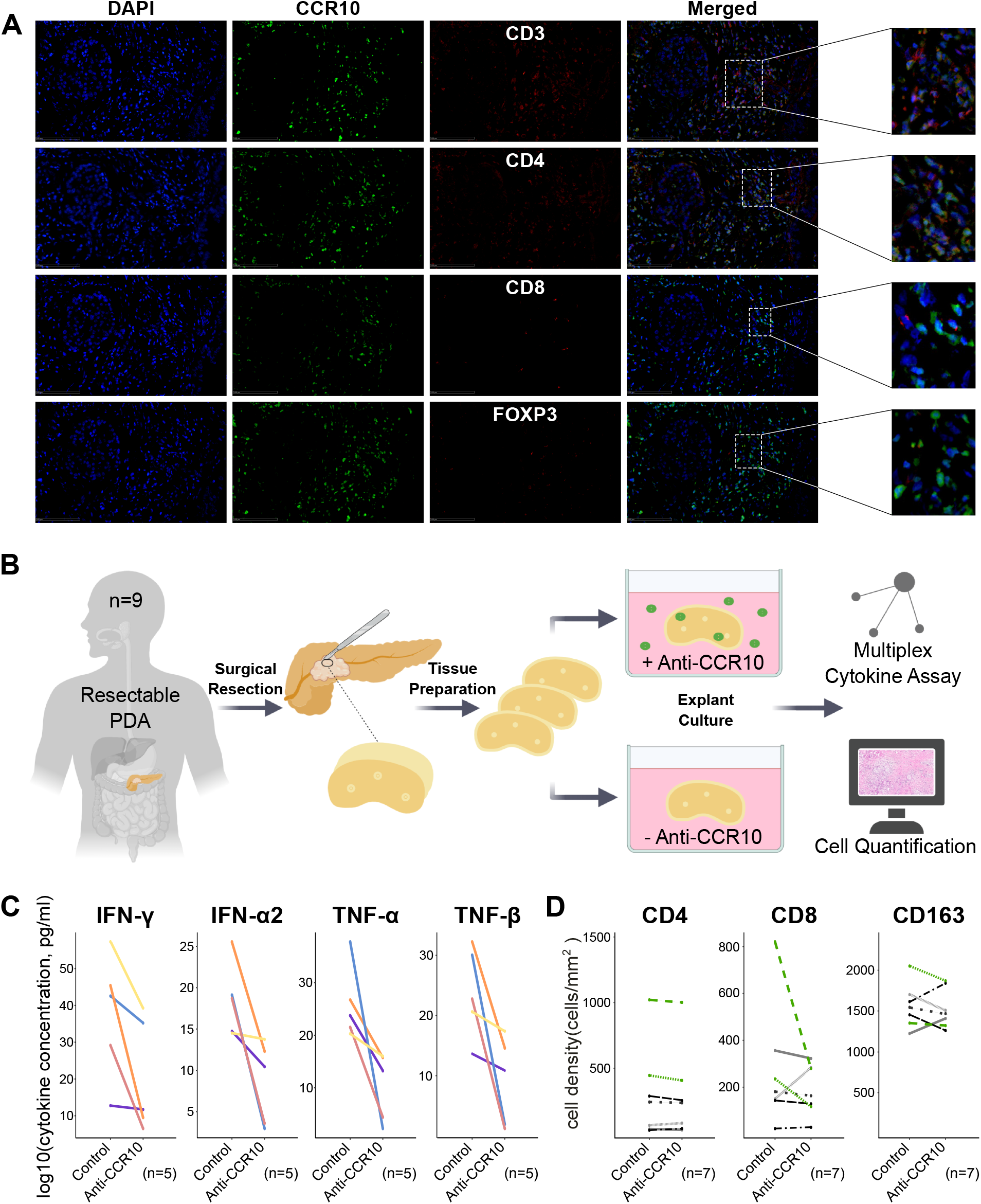
CCR10 is expressed on CD4+ FoxP3-T cells and selective inhibition of CCR10 abrogates the TH1-type cytokine profile **(A)** Immunofluorescence of serial human PDA tissue samples as indicated. Scale bars, 100 μm. **(B)** Schematic overview of workflow as indicated. **(C)** Cytokine alterations within the explant model after 24 hr using human PDA tissue samples. Data from five different patients is presented before and after treatment with the small molecule inhibitor of CCR10 (Anti-CCR10). **(D)** Alteration of the intratumoral CD4+, CD8+ and CD163+ immune cell infiltration within the explant model using human PDA tissue samples. Data from seven different patients is presented before and after treatment with the small molecule inhibitor of CCR10 (Anti-CCR10). The two tissues with the highest absolute number of intratumoral CD4+ cells are indicated in green. PDA= pancreatic ductal adenocarcinoma See also Figure S3.

To assess whether blockade of CCR10 abolishes anti-tumor properties in the microenvironment of PDA, we investigated the effect of a selective small molecule inhibitor of CCR10 (BI-6901) in a patient-based organotypic tumor explant model^20^ **(Table S2) (Figure 3B)**. Inhibition of CCR10 led to an abrogation of the T_H_1-type cytokine program by downregulating the expression of IFN-γ, IFN-α2, TNF-α and TNF-β **(Figure 3C)**. Other T_H_1-supporting cytokines (TRAIL, IL-12p40, IL-12p70, IL-2, IL-7, IL-10) showed the same alteration **(Figure S3A)**. At the cellular level no difference was observed in the quantity of CD163^+^ or CD4^+^ cells upon blockade of CCR10 **(Figure 3D)**. However, the two tissues with the highest absolute number of CD4^+^ lymphocytes showed the clearest reduction in tumor-infiltrating cytotoxic T cells after CCR10 inhibition **(Figure 3D)**.

### Intratumoral Langerhans islet cells secrete CCL27 via STAT3 regulation and simultaneously downregulate insulin production

We aimed to determine the metabolic implications for immunomodulatory CCL27-producing Langerhans islets, which are primarily attributed to the endocrine regulation of glycemic control. To evaluate clinically relevant metabolic features of Langerhans islets, we sought to compare the preoperative glycemic status (type 2 diabetes mellitus (T2DM) or no T2DM) of PDA patients based on their intratumoral CCL27 expression. This analysis unraveled that CCL27 expression is significantly associated with the onset of T2DM in tumor patients **(Figure 4A)**. Furthermore, the marker combination of CCL27 and preoperative diagnosis of diabetes was shown to serve as a potential parameter for favorable prognosis. However, T2DM is a complex disease with multiple negative implications for human health and further investigation is needed to better understand the role of different types of diabetes and the timepoint of disease onset (long-standing vs. new-onset T2DM) in the context of PDA **(Figure 4B) (Table 2)**.

**Figure 4.**
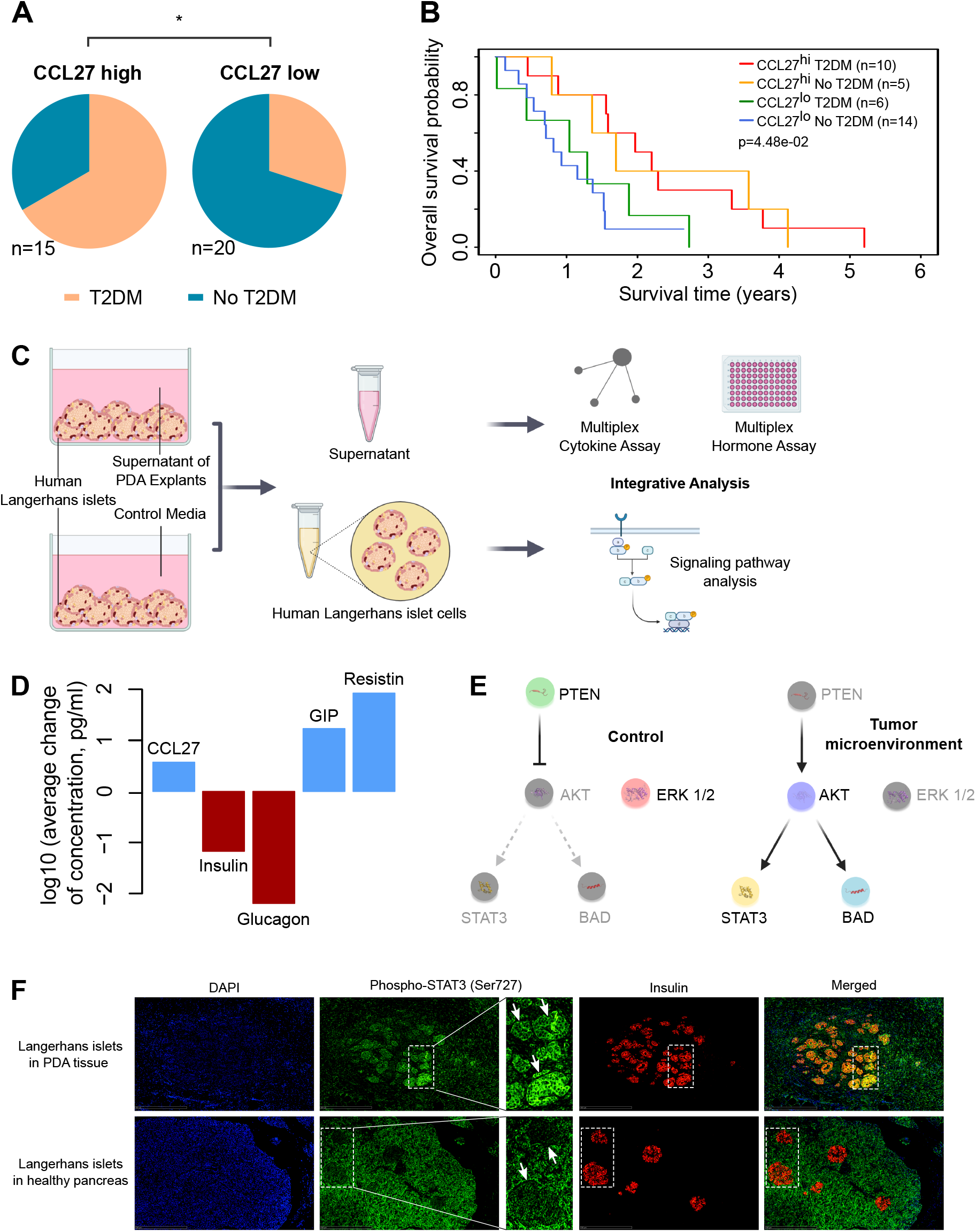
Intratumoral Langerhans islet cells secrete CCL27 via STAT3 regulation and simultaneously downregulate insulin production **(A)** Pie charts showing the distribution of PDA patients with diagnosis of diabetes prior to surgery. The patients were stratified into CCL27 high (hi) (n=15) and CCL27 low (lo) (n=20) group based on their intratumoral CCL27 concentration. **(B)** Kaplan-Meier survival plot of patients based on combination of CCL27 (hi or lo) and T2DM (yes or no). The patients were stratified into CCL27 hi and CCL27 lo group based on their intratumoral CCL27 concentration. The data were analyzed using the log-rank test. **(C)** Schematic overview of workflow as indicated. **(D)** Alteration of indicated protein concentration expressed by human Langerhans islet cells after 48 hr culture in the supernatant of five different human PDA explants (representing five different human pancreatic tumor microenvironments). The expression profile was compared to human Langerhans islet cells cultured in control media for 48 hr. **(E)** Schematic signaling effects of Langerhans islets cells in a pancreatic tumor microenvironment. Grey molecules are inactive and colored molecules are active. **(F)** Immunofluorescence of human PDA tissue and healthy pancreas tissue as indicated. Arrows indicate Langerhans islets. Scale bars, 500 μm. T2DM= type 2 diabetes mellitus PDA= pancreatic ductal adenocarcinoma *p≤0.05. See also Figure S4.

**Table 2.** Subject characteristics

|  |  | CCL27<sup>hi</sup><br>Diabetes | CCL27<sup>hi</sup><br>No diabetes | CCL27<sup>lo</sup><br>Diabetes | CCL27<sup>lo</sup><br>No diabetes | P-value |
| --- | --- | --- | --- | --- | --- | --- |
| Total number of patients (n) |  | 10 | 5 | 6 | 14 |  |
| Female / Male (n / n) |  | 8 / 2 | 3 / 2 | 1 / 5 | 4 / 10 | 0.033 |
| Age | Mean (years) | 68.9±0.9 | 62.2±1.7 | 65.7±1.7 | 61.8±0.9 |  |
|  | > / ≤ Average age<sup>a</sup><br>(n / n) | 8 / 2 | 2 / 3 | 3 / 3 | 7 / 7 | 0.353 |
| BMI<sup>b</sup> | Mean (kg/m<sup>2</sup>) | 25.7±0.5 | 21.8±0.7 | 23.1±0.5 | 24.7±0.2 |  |
|  | > / ≤ Average BMI<sup>c</sup><br>(n / n) | 6 / 2 | 1 / 4 | 1 / 4 | 6 / 4 | 0.153 |
| T | T3 (n) | 10 | 5 | 6 | 13 | 1.000 |
|  | T4 (n) | 0 | 0 | 0 | 1 |  |
| N | N0 (n) | 3 | 0 | 0 | 2 | 0.503 |
|  | N1 (n) | 7 | 5 | 6 | 12 |  |
| M | M0 (n) | 10 | 5 | 6 | 13 | 1.000 |
|  | M1 (n) | 0 | 0 | 0 | 1 |  |
| Grade | G2 (n) | 6 | 2 | 5 | 9 | 0.592 |
|  | G3 (n) | 4 | 3 | 1 | 5 |  |
| R<sup>d</sup> | R0 (n) | 2 | 0 | 0 | 2 | 0.731 |
|  | R1 (n) | 8 | 5 | 6 | 11 |  |
| HbA1c<br>level<sup>e</sup> | Mean (%) | 7.0±0.08 | 5.8±0.06 | 7.5±0.20 | 5.4±0.04 |  |
|  | > / ≤ Average<br>HbA1c<sup>f</sup> (n / n) | 9 / 1 | 1 / 4 | 6 / 0 | 0 / 13 | < 0.0001 |
| Gemcitabine-based<br>chemotherapy (n)<sup>g</sup> |  | 10 | 4 | 3 | 10 |  |
| Overall survival (days) |  | 849±52 | 843±106 | 450±60 | 370±17 |  |

Data are shown as mean ± SEM.
Fisher's exact test was used to compare patients of the different groups.
T= stage of primary tumor, N= regional lymph node status, M= distant metastasis status, R= resection margin status, BMI= Body-Mass-Index.
<sup>a</sup> Average age in all patients is 64.5 years.
<sup>b</sup> Information available on BMI from 28 patients.
<sup>c</sup> Average BMI in all patients is 24.18 kg/m<sup>2</sup>.
<sup>d</sup> Information available on resection margin status from 34 patients.
<sup>e</sup> Information available on HbA1c level from 34 patients.
<sup>f</sup> Average HbA1c level in all patients is 6.3 %.
<sup>g</sup> Standard treatment in the study period was upfront resection and gemcitabine-based adjuvant chemotherapy. Information available on confirmed application of gemcitabine from 27 patients.

To more fully characterize the multifaceted role of Langerhans islets in PDA, we explored their secretion and signaling profile after exerting tumoral stress on them **(Figure 4C)**. The tumor microenvironment for this analysis was mimicked by using the supernatant of PDA explants from five different patients. We observed that tumoral stress led to an increased secretion of CCL27 matching our previous findings **(Figure 4D)**. Simultaneously, the secretion of insulin and glucagon was downregulated **(Figure 4D)**. While on the other hand, the secretion of the incretin glucose-dependent insulinotropic polypeptide (GIP) as well as the production of the hormone resistin was increased in Langerhans islet cells **(Figure 4D)**. The same tendency (upregulation in 3 of 5 cases) was seen for glucagon-like peptide 1 (GLP-1) **(Figure S4A)**. However, evaluation of dynamic changes in hormone concentrations is necessary to comprehensively understand these regulatory mechanisms.

Molecular analyses based on protein phosphorylation patterns in these Langerhans islets cells showed an activation of signal transducer and activator of transcription 3 (STAT3) and BCL2-associated agonist of cell death (BAD) upstream regulated by phosphatase and tensin homologue (PTEN) and AKT (also known as protein kinase B or PKB). Simultaneously, the extracellular signal-regulated kinases 1 and 2 (ERK1/2) were dephosphorylated and thereby inactivated **(Figure 4E and S4B)**. Further, we corroborated the phosphorylation of STAT3 in intratumoral Langerhans islets via immunofluorescence staining by comparing human PDA tissue and healthy pancreas tissue **(Figure 4F)**.

### T2DM and dynamic of HbA1c levels are markers for response to neoadjuvant chemotherapy

To explore the relationship of tumoral CCL27 expression and treatment response to adjuvant chemotherapy (= number of adjuvant chemotherapy cycles), we performed a comparison analysis between tissues of patients with high tumoral CCL27 expression (CCL27^hi^) and low tumoral CCL27 expression (CCL27^lo^). We discovered that CCL27 expression, also in a linear relationship, was significantly associated with the number of chemotherapy cycles **(Figure 5A)**.

**Figure 5.**
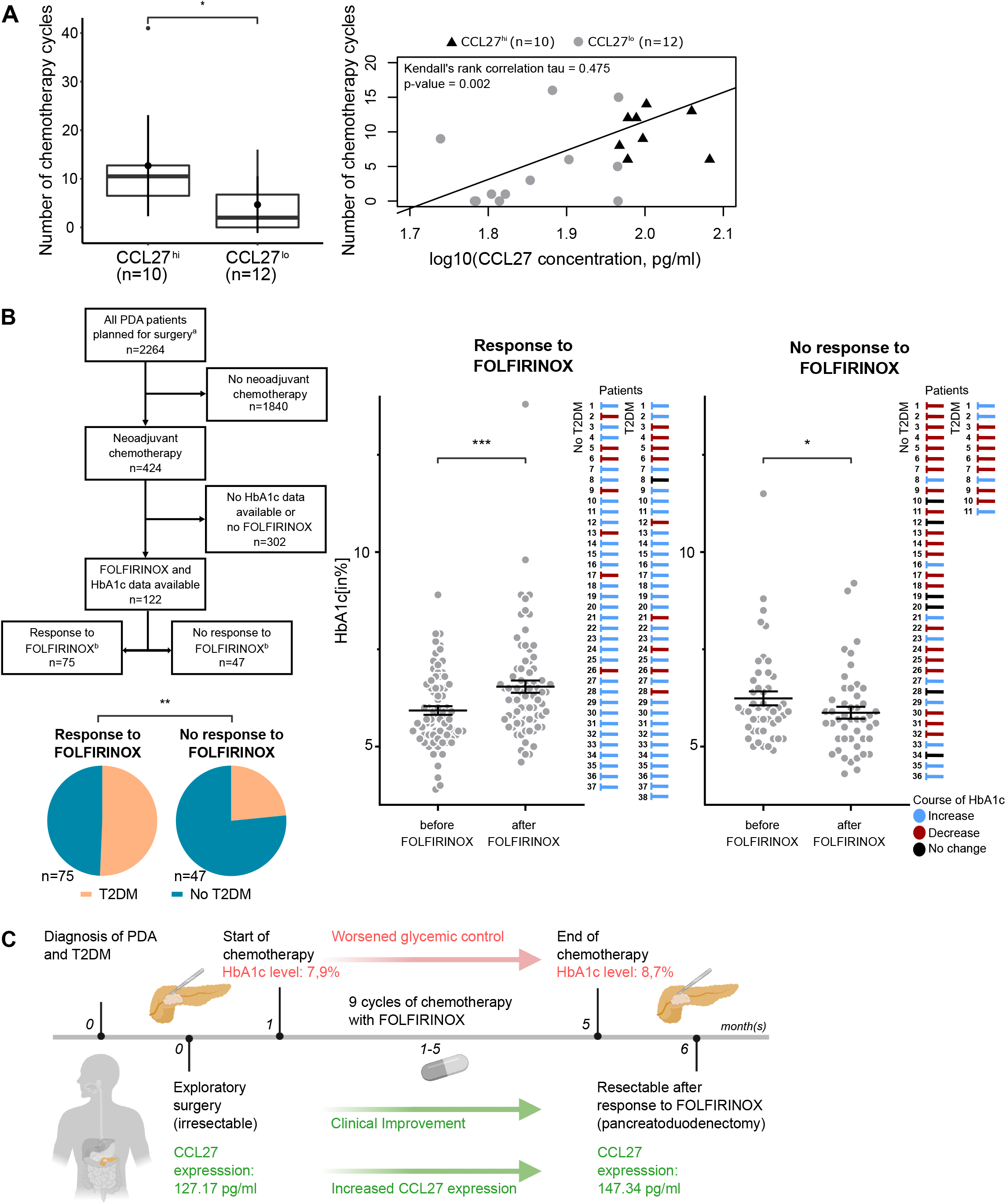
T2DM and dynamic of HbA1c levels are markers for response to neoadjuvant chemotherapy **(A)** Comparison between CCL27 high (hi) and CCL27 low (lo) group based on the number of chemotherapy cycles. Standard treatment regimen was FOLFIRINOX as neoadjuvant chemotherapy. The patients were stratified into CCL27 hi and CCL27 lo group based on their intratumoral CCL27 concentration (left). Correlation analysis showing the association of CCL27 concentrations with number of chemotherapy cycles. The analysis was performed using the Kendall rank correlation (right). **(B)** Retrospective analysis performed on a total of 2264 patients. HbA1c levels (before and after FOLFIRINOX) were available from 122 patients who were treated with neoadjuvant chemotherapy. The pie charts are showing the distribution of PDA patients with diagnosis of T2DM at the beginning of neoadjuvant chemotherapy with FOLFIRINOX. The patients were stratified into Response to FOLFIRINOX (n=75) and No response to FOLFIRINOX (n=47). Comparison of HbA1c levels before and after FOLFIRINOX treatment in responders and non-responders. The course of HbA1c levels of individual patients are described as indicated (blue=increase, red=decrease, black=no change). **(C)** Timeline describing the case of a patient as indicated. *p≤0.05, **p≤0.005, ***p≤0.0001. a All PDA patients planned for pancreatoduodenectomy from 2016-2020 in the University Hospital Heidelberg were included. b Response to FOLFIRINOX was assumed when the tumor was resectable. No response to FOLFIRNOX was assumed when the tumor remained irresectable. Data are represented as mean ± SEM and compared by two-sided Student’s t test. PDA= pancreatic ductal adenocarcinoma T2DM= type 2 diabetes mellitus FOLFIRINOX= fluorouracil, leucovorin, irinotecan and oxaliplatin See also Figure S5.

Since CCL27 secretion is linked to glycemic control, we hypothesized that diagnosis of T2DM or glycated haemoglobin (HbA1c) levels may have the potential to identify patients who are most likely to respond to chemotherapy. To investigate this, we screened a cohort of 2264 PDA patients who were planned for pancreatoduodenectomy **(Figure B)**. We identified 122 patients who had undergone neoadjuvant chemotherapy with FOLFIRINOX prior to the planned surgery and with extractable data on HbA1c levels before and after FOLFIRINOX **(Table 3)**. The retrospective analysis unveiled that diagnosis of T2DM was significantly associated with response to neoadjuvant chemotherapy with FOLFIRINOX **(Figure 5B)**. Moreover, decline of glycemic control (= increase of HbA1c levels) during neoadjuvant chemotherapy with FOLFIRINOX was also significantly associated with treatment response. This observation was statistically significant in PDA patients with and without T2DM **(Figure S5A)**. In contrast, improved glycemic control (= decrease of HbA1c levels) was significantly associated with unresponsiveness to neoadjuvant FOLFIRINOX **(Figure 5 B)**. This was significant in patients without T2DM and the same trend was seen for patients with T2DM **(Figure S5A)**.

**Table 3.**
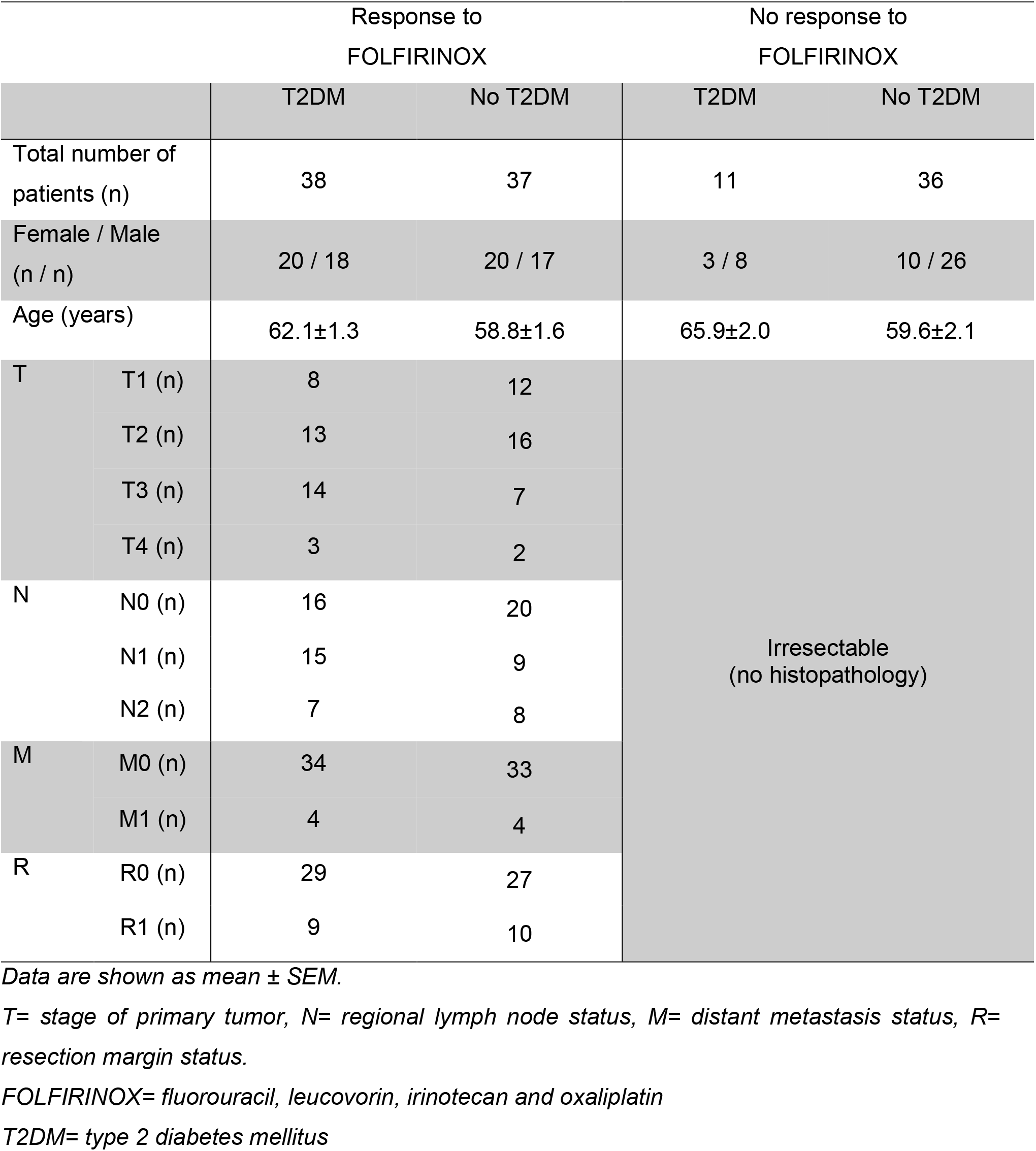
Subject characteristics

To confirm our concept that deterioration of glycemic control is accompanied by increased local CCL27 levels, we analyzed the tumor tissue of a patient who responded to neoadjuvant chemotherapy with FOLFIRINOX and had a rising HbA1c level **(Figure 5C)**. This case validated our findings and demonstrated that not only did the HbA1c level increase, but also the local CCL27 concentration **(Figure 5 C and S5B)**.

## DISCUSSION

Induction of anti-tumor immunity by increasing the number of tumor-infiltrating T cells is an unresolved challenge and limited T cell infiltration is a major reason for resistance to immunotherapy in PDA. In the present report, we demonstrate a crucial role of endocrine Langerhans islets in shaping anti-tumor immunity by undergoing a phenotype shift from glycemic control to CCL27 production. Subsequently, CCL27 promotes a T_H_1-type cytokine program in the microenvironment, enhances tumoral T cell infiltration and improves survival of patients. To date, CCL27 has primarily been described as a chemotactic mediator of T cell-dependent (skin) inflammation and previous reports showed concordantly that CCL27 induces T cell attraction and inflammation via CCR10^+^ CD4^+^ T lymphocytes whereas CD8^+^ T lymphocytes show negligible levels of CCR10 on their cell surface^29,30^. Our data corroborated these findings in the context of PDA and demonstrated that the anti-tumor effect can be abrogated by inhibition of CCR10. In other malignancies CCL27-CCR10 interaction has been suspected to promote lymph node metastases^31^ and contribute to inflammation-driven hepatocarcinogenesis^32^. However, our results highlight that CCL27-CCR10 interaction plays an opposite role in the microenvironment of PDA by potentially turning it into an immunologically “hot” tumor with beneficial clinical outcomes. Also, our data indicate that efficacy of effector T cell-attraction by CCL27 is dependent on the quantity of CCR10^+^ CD4^+^ T cells, in order to generate a sufficient T_H_1-cytokine signal. This is line with existing evidence, which showed that CD4^+^ T cells play a fundamental role in driving anti-tumor CD8^+^ T cell responses and that their presence is associated with the number of CD8^+^ T cells in solid tumors^33,34^. The use of a fully human patient-derived model in our experiments allowed us to link the clinical observations to functional molecular findings.

Another interesting aspect of our results is that endocrine Langerhans islet cells were identified as the primary source of CCL27 suggesting a critical involvement of the endocrine-exocrine axis in anti-tumor immunity. Primarily, Langerhans islets are attributed to the endocrine regulation of glycemic control, which requires a precisely fine-tuned balance of multiple glucoregulatory hormones. Glucose homeostasis is primarily regulated by a tug-of-war between endocrine Langerhans islet hormones. Glucagon is secreted by α-cells and increases plasma glucose levels, whereas insulin from β-cells decreases them. Furthermore, a variety of other hormones are critically involved, such as the incretins GIP and GLP-1, which are secreted by α-cells and regulate blood glucose levels through potentiation of insulin secretion and inhibition of glucagon secretion^35,39^. In light of the existing evidence, our findings suggest that while intratumoral β-cells secrete CCL27 which inhibits tumor progression, their metabolic function declines resulting in an insulin deficiency. This eventually leads to a compensatory upregulation of intrainsular GIP and GLP-1 secretion, which fail to properly potentiate insulin expression but inhibit glucagon secretion. This mechanism of endocrine-exocrine interaction also explains our clinical observation that onset of T2DM in PDA patients is significantly associated with CCL27 expression of tumors. In addition, the observed lower insulin levels might further help to limit aberrant activation of oncogenic signaling pathways and may contribute to systemic antineoplastic functions^40-42^. Another factor potentially contributing to the loss of glycemic control in patients is the upregulated resistin secretion. Resistin is secreted by adipocytes and Langerhans islets, known as a key link between obesity and diabetes and leads to peripheral insulin resistance^43^.

Further, the identification of STAT3 as activator of CCL27 production is in line with previous findings, which presented its common involvement in cytokine release by Langerhans islet cells^44^. Interestingly, STAT3 also plays a critical role in inhibiting insulin secretion in mice^45,46^, which corroborates our findings. And the observed phosphorylation and activation of anti-apoptotic BAD might contribute to preservation of this CCL27-releasing and insulin-inhibiting phenotype of Langerhans islets.

The interplay of T2DM to PDA is complex and, despite numerous studies, not comprehensively understood. However, strong existing evidence demonstrated that (new-onset) T2DM is likely to be a consequence of a tumor microenvironment consisting of progressively growing PDA cells^47^. We complement prior findings by showing that onset of T2DM is a potential sign of functionally shifting Langerhans islets towards shaping anti-tumor immunity via CCL27 production as a defense mechanism against PDA. This observation also underscores the complex and bilateral role of local immunity in PDA: While IL-1β-induced pancreatitis promotes PDA via immunosuppression^48^, development of T2DM promotes autoimmune anti-tumor responses via CCL27. Further, the interconnection of glycemic control and PDA prompted us to investigate metabolic parameters as markers for response to neoadjuvant chemotherapy with FOLFIRINOX. Historically, the combination therapies FOLFIRINOX and gemcitabine with nab-paclitaxel significantly increased survival of patients^49,50^. While FOLFIRINOX is more effective than gemcitabine, the regimen also causes more side effects and significantly reduces quality of life of patients. However, response to chemotherapy varies, with only one third of patients responding to a specific regimen^51^. Given the fact that patients with unresponsiveness to neoadjuvant chemotherapy and irresectable PDA have a median life expectancy of less than a year preventing FOLFIRINOX-induced toxicity is an urgent clinical need. FOLFIRINOX is known to induce an anti-tumor immune infiltrate in the pancreatic tumor microenvironment, which is characterized by increased cell densities of CD8^+^ and CD4^+^ T lymphocytes as well as decreased numbers of FoxP3^+^ and CD163^+^ cells^52-54^. We hypothesize that this local immune reaction is the reason for our observation of elevated HbA1c levels and increased CCL27 secretion after exposure to FOLFIRINOX, similarly to the production of inflammatory signals as an Langerhans islet response to various local inflammatory factors during development of diabetes^14,55,56^. Our findings highlight that the presence of T2DM before the start of neoadjuvant chemotherapy as well as the dynamic of HbA1c levels during the treatment are markers for response to FOLFIRINOX. In line with this, another independent clinical investigation demonstrated that elevated HbA1c levels can be utilized as a marker for stratifying patients most likely to respond to FOLFIRINOX^51^.

In conclusion, our findings describe a novel mechanism of anti-tumor action in PDA unravelling an unexpected role for endocrine Langerhans islets with implications for the clinical management of PDA. Repurposing metabolic parameters has the potential to serve as a tool for clinical decision-making. However, prospective clinical trials are needed to further investigate these applications. Also, CCL27-CCR10 interaction could serve as a stratification parameter for immunotherapeutic strategies.

## ACKNOWLEDGEMENTS

We thank Jana Wolf, Ulrike Prüfer and Rosa Eurich for excellent technical assistance. Schematic figures were created with BioRender.com. PDA samples were provided by the Pancobank platform at the European Pancreas Center Heidelberg (EPZ), member of the Biomaterial Bank Heidelberg (BMBH).

## CONTRIBUTORS

Conceptualization, A.A. and N.H.; Methodology, A.A., S.K., M.S. and N.H.; Formal Analysis, A.A., P.C., D.F. and N.H.; Investigation, A.A., S.K. and R.K.; Resources, R.K., T.H., N.G., M.H., C.S., C.F., I.Z., D.J. and N.H.; Data Curation, A.A., P.C. and N.H.; Writing - Original Draft, A.A.; Writing - Review & Editing, A.A, R.K., S.K., M.S, N.V., D.F., M.H., N.G., C.S., C.F., M.S., T.S., Y.T., L.Z., I.Z., D.J. and N.H.; Visualization, A.A., P.C., N.V. and N.H.; Supervision, I.Z., D.J. and N.H.; Project Administration, A.A. and N.H.; Funding Acquisition, I.Z., D.J. and N.H.

## ETHICS APPROVAL

Ethics Committee of the University of Heidelberg.

## DATA AVAILABILITY STATEMENT

All data are available upon request. All data relevant to the study are included in the article or uploaded as supplementary information.

