## Supplemental information for "Langerhans islets induce anti-tumor immunity at the expense of glycemic control and predict chemotherapy response in pancreatic cancer"

#### SUPPLEMENTAL TABLES

Table S1. Subject characteristics (Cohort 1)

| | | Cohort 1 |
|---|---|---|
| Total number of patients (n) | | 51 |
| Female / Male (n / n) | | 25 / 26 |
| Age (years) | | 65.6±1.4 |
| T | T3 (n) | 50 |
| | T4 (n) | 1 |
| N | N0 (n) | 8 |
| | N1 (n) | 43 |
| M | M0 (n) | 48 |
| | M1 (n) | 3 |
| Grade | G2 (n) | 33 |
| | G3 (n) | 18 |
| R[a] | R0 (n) | 6 |
| | R1 (n) | 44 |
| Diabetes prior to surgery (yes (n) / no (n))[b] | | 16 / 19 |
| Overall survival (days) | | 609±68 |

Data are shown as mean ± SEM.

T= stage of primary tumor, N= regional lymph node status, M= distant metastasis status, R= resection margin status.

[a] Information available on resection margin status from 50 patients.

[b] Information available on glycemic status (according to HbA1c levels or medical history) from 35 patients.

Table S2. Subject characteristics (Cohort 2)

| | | Cohort 2 |
|---|---|---|
| Total number of patients (n) | | 9 |
| Female / Male (n / n) | | 4 / 5 |
| Age (years) | | 66.2±2.9 |
| T | T3 (n) | 8 |
| | T4 (n) | 1 |
| N | N0 (n) | 4 |
| | N1 (n) | 3 |
| | N2 (n) | 2 |
| M | M0 (n) | 8 |
| | M1 (n) | 1 |
| Grade[a] | G2 (n) | 3 |
| | G3 (n) | 3 |
| R | R0 (n) | 5 |
| | R1 (n) | 3 |
| Neoadjuvant therapy with FOLFIRINOX (n) | | 3 |

Data are shown as mean ± SEM.

T= stage of primary tumor, N= regional lymph node status, M= distant metastases status, R= resection margin status, FOLFIRINOX= fluorouracil, leucovorin, irinotecan, oxaliplatin.

[a] Grading is not performed in patients with neoadjuvant chemotherapy.

Figure S1

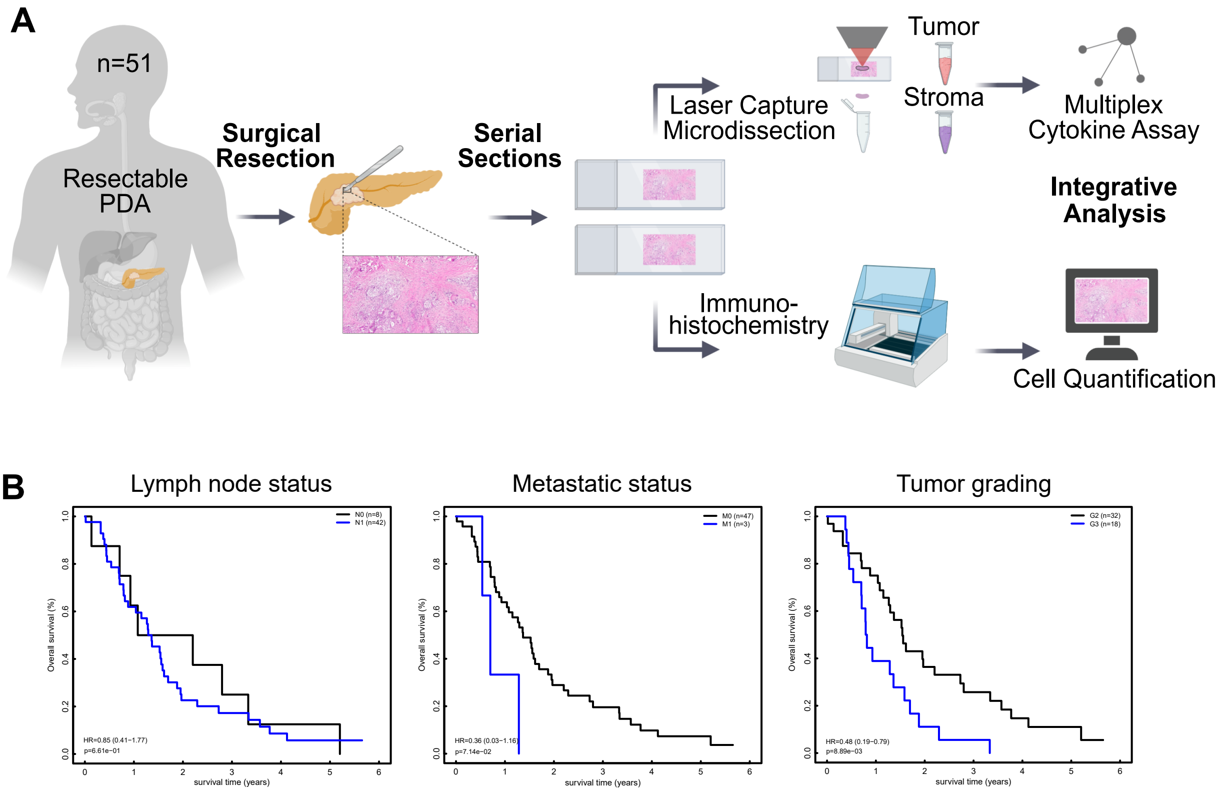
A
n=51
Resectable PDA
Surgical Resection
Serial Sections
Laser Capture Microdissection
Tumor
Stroma
Multiplex Cytokine Assay
Integrative Analysis
Immuno-histochemistry
Cell Quantification
B
Lymph node status
N0 (n=8)
N1 (n=42)
HR=0.85 (0.41−1.77)
p=6.61e−01
Metastatic status
M0 (n=47)
M1 (n=3)
HR=0.36 (0.03−1.16)
p=7.14e−02
Tumor grading
G2 (n=32)
G3 (n=18)
HR=0.48 (0.19−0.79)
p=8.89e−03
Overall survival (%)
survival time (years)

### Figure S3

**A**

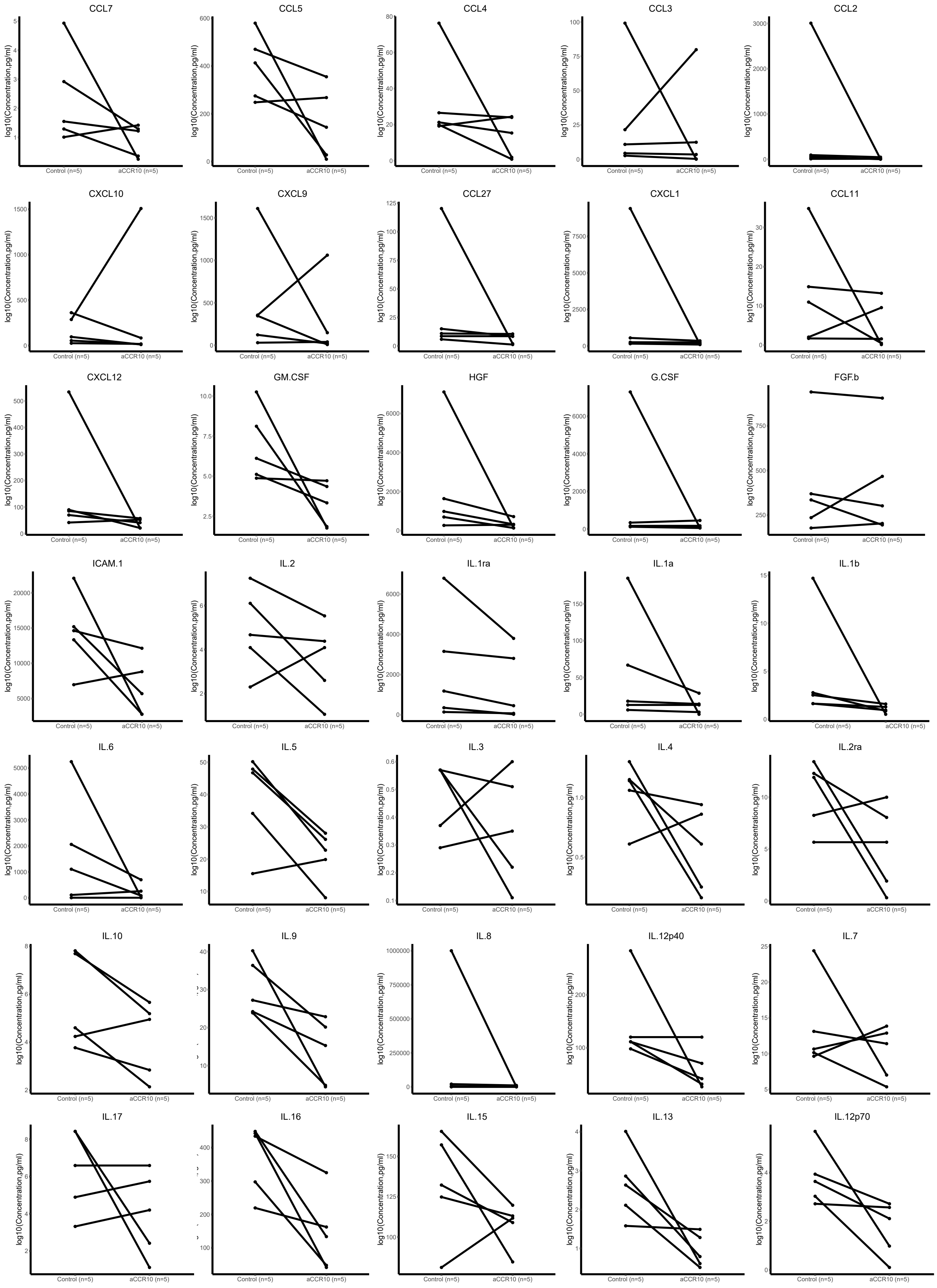

CCL7
CCL5
CCL4
CCL3
CCL2
CXCL10
CXCL9
CCL27
CXCL1
CCL11
CXCL12
GM.CSF
HGF
G.CSF
FGF.b
ICAM.1
IL.2
IL.1ra
IL.1a
IL.1b
IL.6
IL.5
IL.3
IL.4
IL.2ra
IL.10
IL.9
IL.8
IL.12p40
IL.7
IL.17
IL.16
IL.15
IL.13
IL.12p70
log10(Concentration,pg/ml)
Control (n=5)
aCCR10 (n=5)

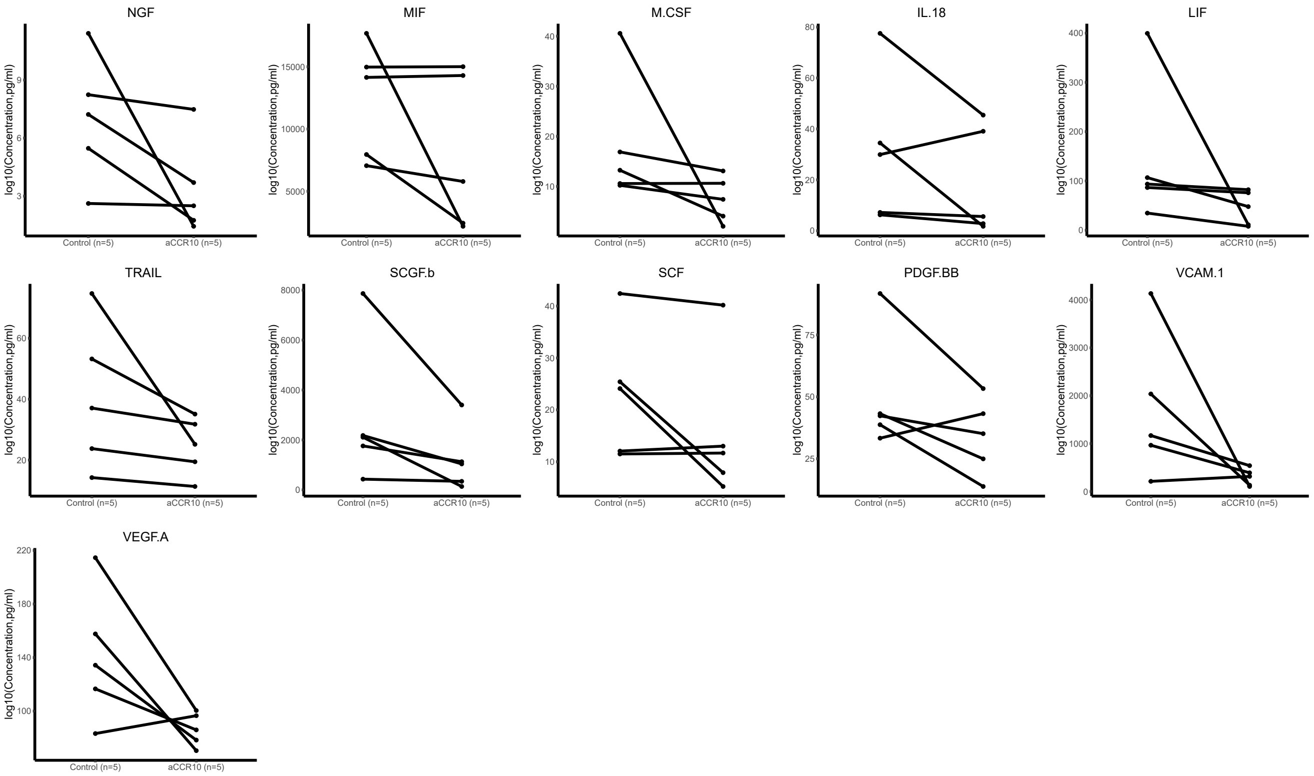
NGF
MIF
M.CSF
IL.18
LIF
TRAIL
SCGF.b
SCF
PDGF.BB
VCAM.1
VEGF.A
log10(Concentration,pg/ml)
Control (n=5)
aCCR10 (n=5)

Figure S4

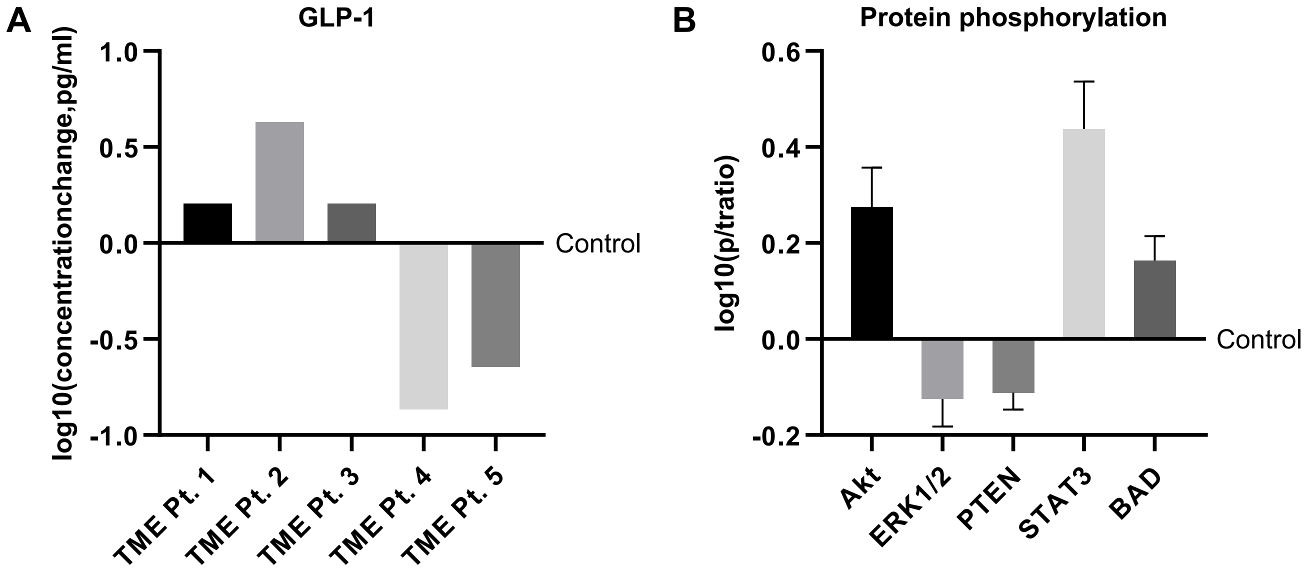
A
GLP-1
log10(concentrationchange,pg/ml)
1.0
0.5
0.0
-0.5
-1.0
Control
TME Pt. 1
TME Pt. 2
TME Pt. 3
TME Pt. 4
TME Pt. 5
B
Protein phosphorylation
log10(p/tratio)
0.6
0.4
0.2
0.0
-0.2
Control
Akt
ERK1/2
PTEN
STAT3
BAD

Figure S5

A

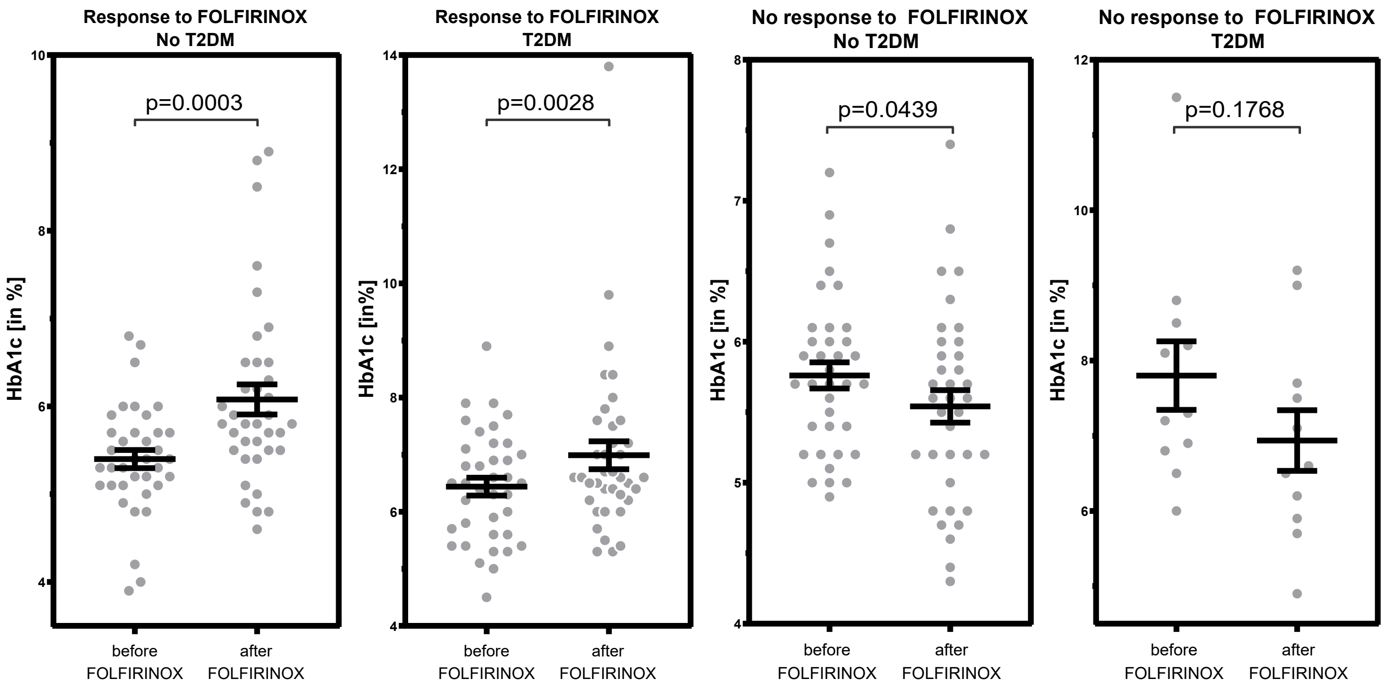

Response to FOLFIRINOX No T2DM
p=0.0003
HbA1c [in %]
before FOLFIRINOX
after FOLFIRINOX
Response to FOLFIRINOX T2DM
p=0.0028
HbA1c [in%]
before FOLFIRINOX
after FOLFIRINOX
No response to FOLFIRINOX No T2DM
p=0.0439
HbA1c [in %]
before FOLFIRINOX
after FOLFIRINOX
No response to FOLFIRINOX T2DM
p=0.1768
HbA1c [in %]
before FOLFIRINOX
after FOLFIRINOX


B

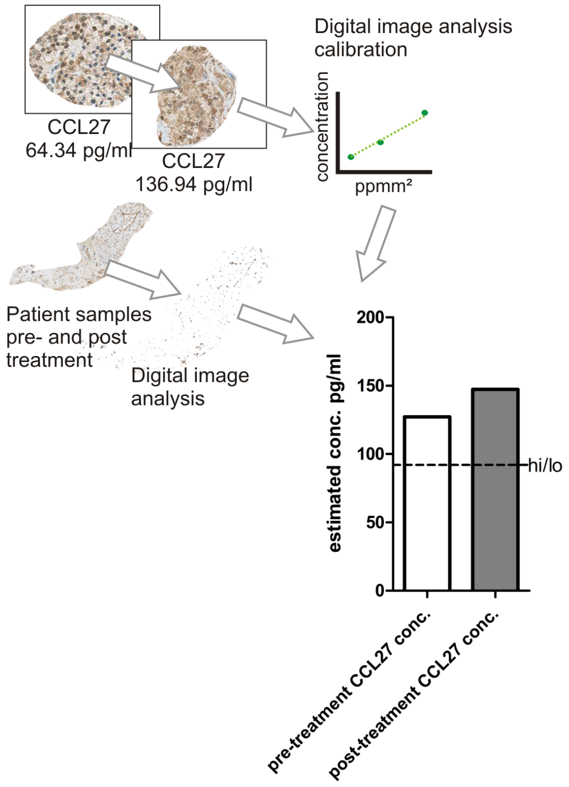

CCL27
64.34 pg/ml
CCL27
136.94 pg/ml
Digital image analysis calibration
concentration
ppmm²
Patient samples pre- and post treatment
Digital image analysis
estimated conc. pg/ml
200
150
100
50
0
hi/lo
pre-treatment CCL27 conc.
post-treatment CCL27 conc.

#### SUPPLEMENTAL FIGURE LEGENDS

**Figure S1** Intratumoral CCL27 expression is associated with improved overall survival

**(A)** Schematic overview of workflow as indicated.

**(B)** Kaplan-Meier survival plots of patients (n=50) based on clinical information as indicated. Survival data were analysed using the log-rank test.

**Figure S3** CCR10 is expressed on CD4+ FoxP3$^-$ T cells and selective inhibition of CCR10 abrogates the $T_H$1-type cytokine profile

**(A)** Cytokine alterations within the explant model after 24 hr using human pancreatic cancer tissue samples. Data from five different patients are presented before and after treatment with the small molecule inhibitor of CCR10 (Anti-CCR10).

**Figure S4** Intratumoral Langerhans islet cells secrete CCL27 via STAT3 regulation and simultaneously downregulate insulin production

**(A)** Alteration of GLP-1 levels expressed by human Langerhans islet cells after 48 hr culture in the supernatant of five different human pancreatic cancer explants (representing five different human pancreatic tumor microenvironments (TME)). The expression profile was compared to human Langerhans islet cells cultured in control media for 48 hr.

**(B)** Changes in phosphoprotein levels (phosphoprotein to total protein levels, as described previously(1)) in human Langerhans islet cells within pancreatic tumor microenvironments (mimicked by the supernatant of pancreatic cancer explants of five different patients). Their levels were compared to human Langerhans islet cells cultured in control media.

**Figure S5** CCL27 expression and HbA1c levels are potential markers for response to chemotherapy

**(A)** Comparison of HbA1c levels before and after FOLFIRINOX treatment in responders and non-responders based on diagnosis of T2DM (T2DM or no T2DM).

**(B)** Schematic overview of CCL27 quantification using stained tissue sections.

Data are represented as mean ± SEM and compared by two-sided Student’s t test.
